# Boundary sequences flanking the mouse tyrosinase locus ensure faithful pattern of gene expression

**DOI:** 10.1101/2020.06.07.139048

**Authors:** Davide Seruggia, Almudena Fernández, Marta Cantero, Ana Fernández-Miñán, José Luis Gomez-Skarmeta, Pawel Pelczar, Lluis Montoliu

## Abstract

Control of gene expression is dictated by cell-type specific regulatory sequences that physically organize the structure of chromatin, including promoters, enhancers and insulators. While promoters and enhancers convey cell-type specific activating signals, insulators prevent the cross-talk of regulatory elements within adjacent loci and safeguard the specificity of action of promoters and enhancers towards their targets in a tissue specific manner. Using the mouse tyrosinase (*Tyr*) locus as an experimental model, a gene whose mutations are associated with albinism, we described the chromatin structure in cells at two distinct transcriptional states. Guided by chromatin structure, through the use of Chromosome Conformation Capture (3C), we identified sequences at the 5’ and 3’ boundaries of this mammalian gene that function as enhancers and insulators. By CRISPR/Cas9-mediated chromosomal deletion, we dissected the functions of these two regulatory elements in vivo in the mouse, at the endogenous chromosomal context, and proved their role as genomic insulators, shielding the *Tyr* locus from the expression patterns of adjacent genes.

## INTRODUCTION

Extensive profiling of genome organization and chromatin folding suggested that gene expression and chromatin topology are highly correlated(1). Actively transcribed genes reside in large *A* compartments, mega-base structures characterized by the presence of active chromatin marks and high reciprocal physical interactions(2). Conversely, inactive genetic material is confined to the *B* compartment. Genes can switch their respective compartment(3–5), when they become active during development or differentiation, illustrating the dynamic correlation between gene expression and chromatin topology. At a smaller scale (tens to hundreds of kilobases), highly interacting chromatin regions are organized in topologically associating domains (TADs) characterized by high interdomain interactions(6). TADs partition chromatin in independent units, delimited by boundaries enriched in CTCF-occupied regions. Disruption of boundaries results in misfolding of TADs, with spurious enhancer-promoter interactions that often lead to disease or developmental defects(7–9). Within TADs, enhancers reach their target promoters through intra-TAD chromatin loops, mediated by CTCF, Cohesin, Mediator(10) and other transcription factors(11).

The tyrosinase (*TYR*) gene encodes the rate-limiting enzyme in the biosynthesis of the melanin. Mutations in *TYR* are the molecular cause of the most common type of albinism in Western countries (oculocutaneous albinism type 1, OCA1) a rare disease characterized by severe visual deficits and hypopigmentation in the skin, hair and eyes(12). In the mouse, mutations at *Tyr* affect coat and eye pigmentation, with similar alterations in the visual pathway(13), but without gross detrimental effects on overall physiology and viability. For this reason, *Tyr* has been targeted in mouse(14) and other organisms(15–17) to benchmark transgenesis and genome engineering strategies, from oocyte microinjection of DNA(18, 19) to the most recent CRISPR nuclease(20) and base editor approaches(21, 22). *Tyr* is expressed in neural crest-derived melanocytes populating the skin, the hair bulbs and numerous other body locations, including the choroid, a pigmented layer found between the retina and the sclera of the eye. In addition to melanocytes, *Tyr* is also expressed is the retinal pigmented epithelium (RPE), a monolayer of hexagonal and binucleated pigmented cells located at the outer most layer of the retina, derived from the optic cup(23, 24). Its highly specific expression pattern makes the *Tyr* locus an excellent experimental model locus to explore the mechanisms of gene expression, its relation with chromatin topology and the consequences of genetic perturbations at structurally-relevant sequences. The *Tyr* promoter contains the minimal information necessary to correctly drive *Tyr* expression *in vivo*(18, 25, 26). This transcriptional specificity is conveyed by two highly conserved motifs, the *Tyr Initiator* (*Irn*) and the M-box, both occupied by the transcription factor *Mitf*(27). A regulatory element 5’ upstream in the murine *Tyr* locus was first identified as a DNAse I hypersensitivity site(28). The inclusion of this element in a tyrosinase minigene resulted in copy number-dependant gene expression in transgenic mice(29), suggesting features of a chromatin boundary(30) or a locus control region (LCR)^31,32^. Furthermore, when microinjected into oocytes from albino mice, large yeast artificial chromosomes (YACs) encompassing these *Tyr* upstream sequences fully rescued coat-colour pigmentation(31), whereas shorter constructs, lacking far upstream sequences, provided weak and variegated *Tyr* expression(32, 33). More recently, we deleted the *Tyr* 5’ element at the endogenous locus using a CRISPR/Cas9 approach and generated allelic series to identify a 410 bp core enhancer sequence(34).

Here, using *in vivo* mouse studies, we resolve the chromatin structure of the *Tyr* locus and integrate it with functional data obtained by targeting additional putative cis-regulatory sequences at the 3’ downstream of the *Tyr* gene.

## METHODS

### Cell culture

Mouse melanoma cell line B16-F1, mouse fibroblast NIH-3T3 L929 cells and human embryonic kidney HEK 293 cells were grown in DMEM medium (Dulbecco’s Modified Eagle Medium, Gibco) supplemented with sterile-filtered 10% fetal bovine serum (FBS, Sigma-Aldrich), 2 mM L-glutamine (Invitrogen) and 10 mM HEPES pH 7.4 (Invitrogen) under aseptic conditions using a sterile hood (Telstar Bio II Advance). Cells were cultured at +37°C, 95% of humidity and 5% CO_2_.

### Chromosome conformation capture (3C)

3C analyses were performed as previously reported(35). Briefly, 1×10^7^ B16F0 mouse melanoma or NIH3T3-L929 mouse fibroblast cells were fixed in 2% PFA/PBS. After quenching, cells were resuspended in 500 μl lysis buffer (10 mM Tris HCl, 10 mM NaCl, 0.3% NP40, 1× Roche Complete) and kept on ice. Methyl Green-Pyronin (MGP, Sigma Aldrich) was used to monitor the release of intact nuclei. Chromatin was digested overnight with 300 U *Dpn*II (NEB) at 37C. After enzyme inactivation, chromatin was ligated overnight with 45 Weiss Units of T4 DNA ligase (Promega) in 7 ml ligation buffer (30 mM Tris-HCl, 10 mM MgCl_2_, 10 mM DTT, 1 mM ATP). Finally, the sample was treated with Proteinase K and RNase A. DNA was extracted by phenol-chloroform and resuspended in 150 μl TE (10 mM Tris-HCl, 1 mM EDTA). To generate control libraries as a standard for qPCR, 1 μg of BAC RP24-276I14 and equimolar amount of BAC RP23- 359C16 were mixed, digested with 50 U of *Dpn*I (Roche) and religated with T4 DNA ligase.

### Enhancer blocking assay (EBA) in vitro in mammalian cells

Enhancer-blocking activity was measured as previously reported(36–38) using transient transfection in HEK 293 cells. Putative insulator sequences were cloned in the pELuc vector in between the enhancer and the promoter (*Xho*I, test) and upstream the enhancer (*Pst*I, control). Primer sequences are listed in **Supplementary Table**. Luminescence was measured for each construct in triplicate and used to quantify enhancer-blocking activity. The cHS4 chicken beta-globin insulator and its minimal core motif (II/III) were used as positive control; a mutated version of the II/III sequence was used as negative control. One-way ANOVA with Bonferroni post-hoc correction test was used.

### Enhancer-blocking assay (EBA) in vivo in zebrafish embryos

Assays were carried as reported(37, 39). Putative insulator sequences were cloned in between a hindbrain-specific enhancer and a somite-specific promoter. Single-copy integration in the zebrafish genome was obtained by Tol2-mediated transgenesis in fertilized zebrafish embryos. GFP fluorescence was acquired at 36 hpf and quantified using Laser Pix (BioRad). Enhancer-blocking activity was measured based on relative levels of somite/CNS GFP expression. Statistical analyses (median test) were calculated with IBM-SPSS v.21.

### CRISPR/Cas9-mediated chromosomal deletion

A pair of RNA guides flanking the *Tyr* 3’ downstream element was designed using the crispr.mit.edu online tool and cloned by Golden Gate Cloning into the *Esp*3I sites of MLM3636 plasmid (Addgene #43860). Cas9 mRNA was prepared from hCas9 plasmid (Addgene #41815) as described(34, 40) and injected into B6D2F2 (Harlan; originated from B6CBAF1/OlaHsd) mouse fertilized eggs. Founder animals were bred to albino outbred HsdWin:NMRI (Harlan) mice, the reference albino genetic background used in all previous *Tyr* transgenic mouse studies(29, 32–34). *Tyr* 3’ mouse lines were maintained in albino outbred HsdWin:NMRI background.

Primers used for cloning and genotyping are available in **Supplementary Table.** Whole-mount retinae and Hematoxylin-Eosin stained cortical sections were prepared as described(34). Melanin content of skin and eye biopsies was estimated by optical density(34).

In this study, all genome-edited mouse lines were named as TYRINS3#, followed by a number corresponding to the founder they were derived from, as done for the TYRINS5# lines described before(34). The recommended nomenclature for these newly generated mouse lines is stock-*Ty*^*remXLmon*^, where ‘X’ is the corresponding ordinal number for each genome-edited (endonuclease mediated) mouse line(41). For brevity, TYRINS5# and TYRINS3# mouse gene-edited lines are also referred to as *Tyr* 5’ and *Tyr* 3’, respectively, throughout this manuscript. All animal procedures with mice reported in this work were first validated by the National Centre for Biotechnology Ethics Committee on Animal Experimentation (OEBA), thereafter approved by the National CSIC Ethics Committee and eventually authorized by the Autonomous Community of Madrid, acting as the competent authority, according to the Spanish legislation and the European Directive 2010/63/EU. All mice were housed at the registered CNB animal facility, fed and provided water and regular rodent chow ad libitum, with a light/dark cycle 08:00-20:00, according to the European and Spanish norms, and the animal welfare recommendations. Both male and female individuals were used indistinctly.

### mRNA expression analysis

Total RNA was isolated from cultured cells and tissues using the RNeasy Mini Kit (Qiagen) with on-column DNAse treatment. 500 ng of RNA were retrotranscribed using SuperScript III Reverse Transcriptase (Thermo Fisher). qPCR reactions were performed with TaqMan Universal PCR Master Mix (Thermo Fisher) using probes for *Tyr* (Mm00495817_m1), *Nox4* (Mm00479246_m1), *Grm5* (Mm00690332_m1) and *Tbp* (Mm00446973_m1) as described(42).

## RESULTS

### The Tyr locus is organized into two distinct chromatin loops

The mouse *Tyr* locus spans over 180 kb on chromosome 7 and its expression patter is restricted to melanocytes and in cells of the retinal pigmented epithelium(24). Conversely, genes flanking the *Tyr* locus manifest completely different expression patters as evidenced by the ubiquitous expression of *Nox4*(43) and the central nervous system-specific expression of *Grm5*(44) gene. We used published data(45, 46) to explore the chromatin landscape of the *Tyr* locus. In melanocytes, acetylation at lysine 27 of histone H3 (H3K27ac) highlights the gene promoter and regulatory elements upstream of the gene body. Upstream and downstream *Tyr*, chromatin carries repressive chromatin marks such as tri-methylation at lysine 27 of histone H3 (H3K27me3), covering the genes *Nox4* and *Grm5*, inactive in melanocytes (**Fig 1A**). To explore the chromatin structure of the *Tyr* locus in more details and relate it to known regulatory elements of the locus, we decided to analyze this chromosomal territory by performing chromosome conformation capture (3C) in both *Tyr* expressing cells (B16 melanoma cells) and non-expressing cells (NIH 3T3 L929 fibroblasts). We selected a 367 bp *Dpn*II fragment containing the *Irn* and M-box as anchor fragment and interrogated 3C libraries. The *Tyr* promoter engages in multiple interactions within the *Tyr* locus. A strong interaction was observed within a fragment 80 kb 3’ downstream (thereafter, *Tyr* 3’ element) (**Fig 1B**) and confirmed by Sanger sequencing (**Fig S1B**). The promoter/Tyr3’ interaction is observed in both B16 and L929 cells, indicating the constitutive nature of the chromatin loop, suggesting a structural role, independently of *Tyr* expression. No interactions are detected with control sequences immediately upstream and downstream the *Tyr* 3’ element. A cell-type specific interaction is found ~15 kb 5’ upstream of the *Tyr* gene, co-localizing with the *Tyr* 5’ element, analyzed previously(29, 32, 34), restricted to the *Dpn*II fragment containing the *Tyr* LCR A and B boxes(29, 47) (**Fig 1B**) and confirmed by Sanger sequencing (**Fig S1A**). When this *Tyr* 5’ region was used as bait, we could recover signal only at the *Tyr* promoter confirming the specificity of the interaction. We previously showed that inactivation of this sequence *in vivo* by deletion results in a loss of pigmentation(34), confirming the relevance of such interaction for gene expression. No interaction is observed between the *Tyr* 5’ element and far 5’ upstream elements, including the CNS-2, an element previously described to be involved in *Tyr* expression regulation(48, 49). No contacts are detected between the *Tyr* 5’ and *Tyr* 3’ elements, indicating that these two boundary elements probably compete with the same binding sites in the *Tyr* promoter and are not binding with each other (**Fig 1C**). Finally, the anchor primer was positioned at the CNS-2(48), a sequence observed to be dispensable for *Tyr* expression in transgenic mice generated with different YAC *Tyr* transgenes(32). Using this fragment as bait we could not observe any interaction within the *Tyr* locus in both 3C libraries in the cell lines used (**Fig 1D**).

**Figure 1:**
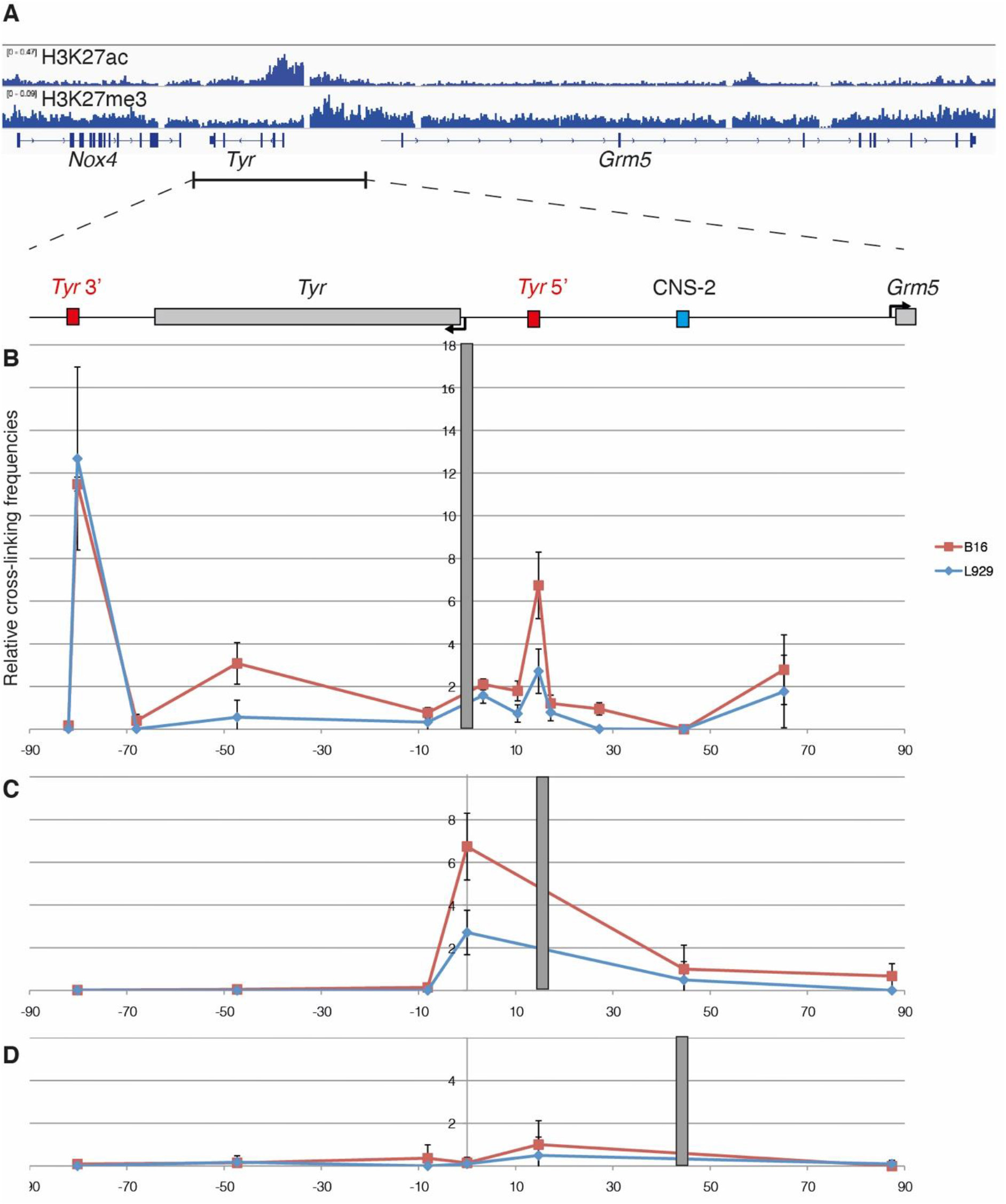
Chromosome conformation of the mouse *Tyr* locus. (**A**) Genomic view of the mouse *Tyr* locus with ChIP-sequencing data obtained from Melan-a cells, including H3K27ac and H3K27me3. Data obtained from Fufa et al. 2015, GEO GSE69950. (**B**) Chromosome Conformation Capture (3C) of mouse B16 melanoma cells (in red) and L292 fibroblasts (in blue). The grey bar indicates the anchor primer on *Tyr* promoter. (**C**) Chromatin loops detected when the anchor is located in correspondence of the *Tyr* 5’ element. (**D**) No interactions are detected when the CNS-2 is used as bait region.

### The Tyr 5’ boundary element contains also a melanocyte-specific enhancer

The anatomical structure of the eye offers the unique opportunity to interrogate the specificity of *Tyr* regulatory elements with regard of the cell of origin. In fact, pigmented, neural-crest (NC) derived melanocytes in the choroid are surrounding the retinal pigmented epithelium (RPE), a single layer of cells at the basis of the neuroretina. We previously demonstrated that skin pigmentation relies on the *Tyr* 5’ element(34). To address the relevance of *Tyr* 5’ in the eye, we compared whole-mount retinae and histological eye sections of wild-type pigmented and *Tyr* 5’ mice. Whole mount retina from *Tyr* 5’ mice display a choroid-specific loss of pigmentation. The typical distribution of hexagonal RPE cells, with unpigmented nuclei, becomes evident in the retina from *Tyr* 5’ mice, whereas is largely masked by underlaying pigmented choroid in the retina from wild-type animals (**Fig. 2A**). Histological analysis confirmed that loss of pigmentation is restricted to the melanocytes present in the choroid, whereas pigmentation of the RPE appears to be unaffected, suggesting that the *Tyr* 5’ element activity is specific to NC-derived cells and dispensable in the RPE (**Fig 2B**). The core sequence(34) of the *Tyr* 5’ enhancer is enriched in Sox10 and Mitf binding motifs (**Fig S2A**), and it is indeed occupied by Sox10 (**Fig 2C**), a NC transcription factor that can activate *Tyr*(50) in melanocytes, but not in RPE, where it is not expressed(51). These results indicates that the *Tyr* 5’ regulatory element, which had been shown to operate as a genomic boundary(30, 34, 39), and be bound by Usf1 factor(48), which has been associated with insulator activity(52), also contains and functions as a transcriptional enhancer(29) to drive *Tyr* expression in neural crest-derived melanocytes in the skin and the choroid, but not in the RPE, in line with previous research that hypothesized the presence of additional RPE-specific regulatory elements(48, 49).

**Figure 2:**
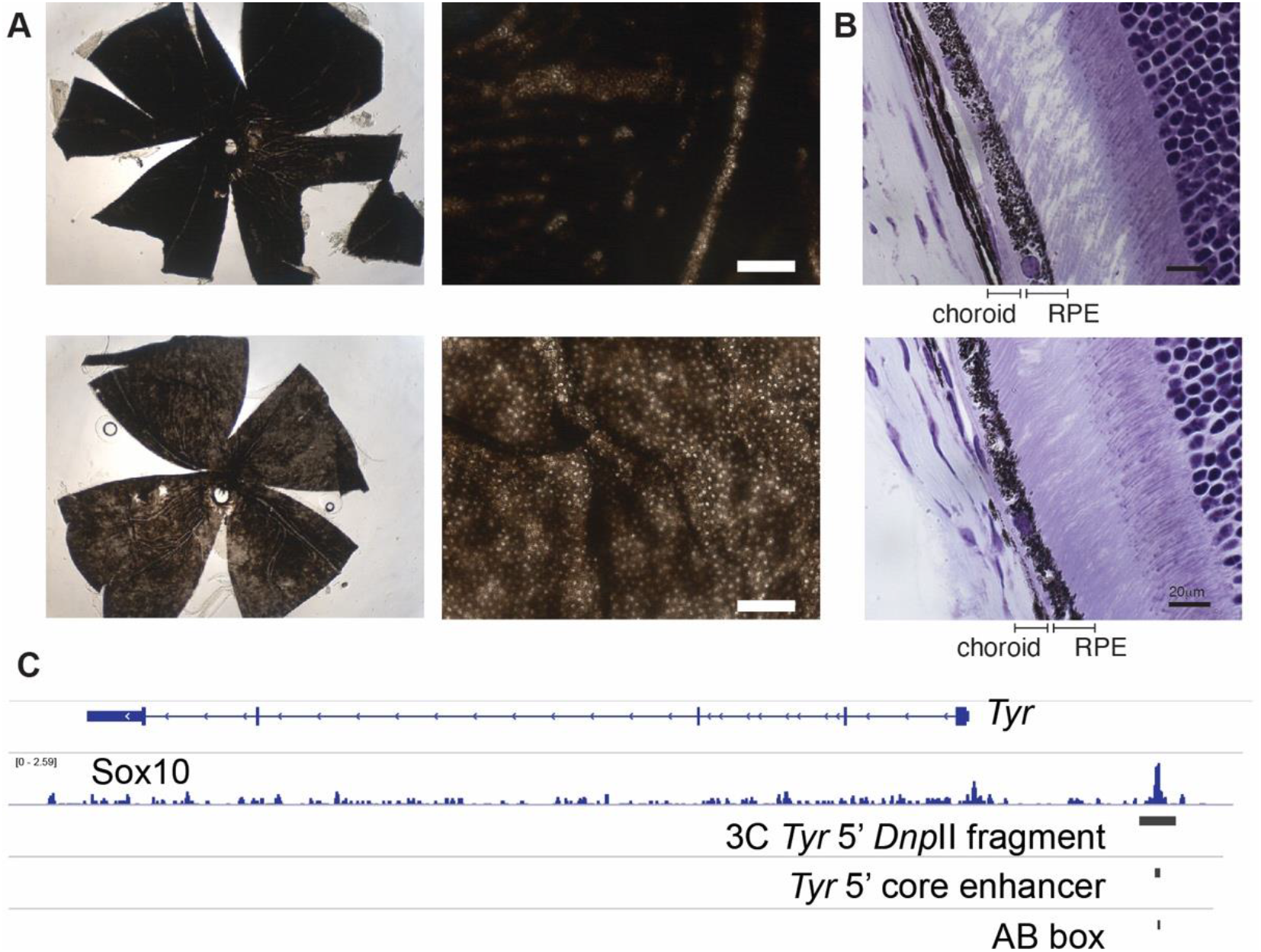
The *Tyr* 5’ element contains a melanocyte-specific *Tyr* enhancer. (**A**) Whole-mount retina from wild-type (top) or TYRINS5 homozygous animals (bottom). (**B**) Hematoxylin-Eosin (HE)-stained coronal sections of eye from adult wild-type (top) or TYRINS5 homozygous mice (bottom). (**C**) Genomic view of the *Tyr* upstream region. A region bound by So×10 in melanocytes(46) overlaps with the DpnII fragment interacting with the *Tyr* promoter, with the core *Tyr* 5’ enhancer and with its AB box.

### The Tyr locus is flanked by chromatin insulators

We previously described and dissected the *Tyr* 5’ element, that coincides with a peak in our 3C assay and whose inactivation *in vivo* results in cell-type specific loss of pigmentation(34). Experiments in transgenic mice and in drosophila(30) and in zebrafish embryos(39) suggested that the *Tyr* 5’ element has insulator activity, in line with previous studies reporting boundary-associated Usf1 factor(52) to bind this *Tyr* 5’ element(48), where its motif is also predicted (**Fig S2A**). Indeed, both the full-length *Tyr* 5’ element and its synthetic GAB core display enhancer-blocking activity in human cells (**Fig 3C**).

**Figure 3:**
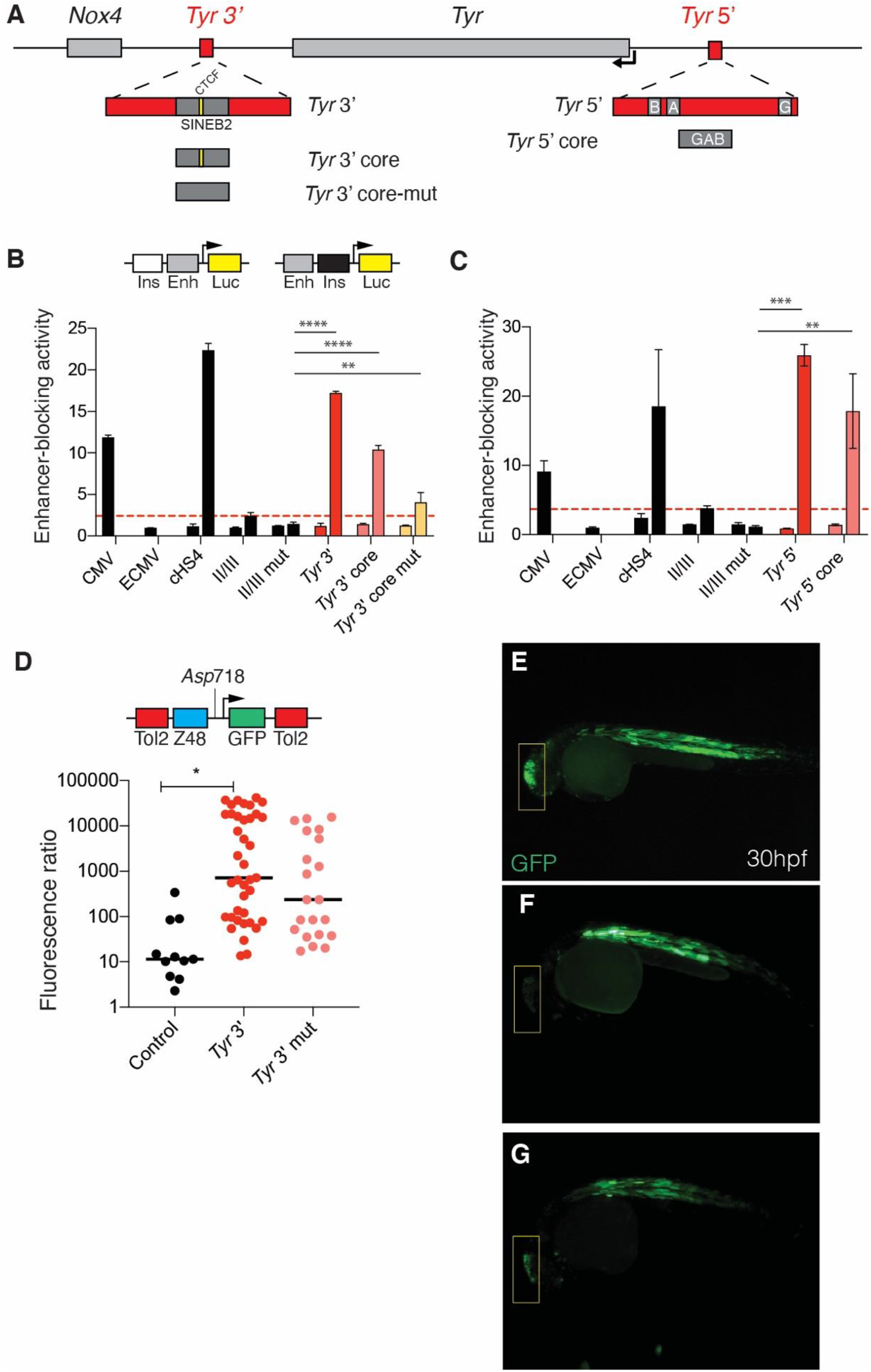
The *Tyr* locus is flanked by chromatin boundaries. (**A**) Schematics of the *Tyr* locus with cartoons illustrating the *Tyr* 3’ and *Tyr* 5’ boundaries. (**B**) Enhancer-blocking assay (EBA) of the *Tyr* 3’ element and its core sequence. Putative insulators are cloned upstream the CMV enhancers (control) or downstream the CMV enhancer (test). Enhancer-blocking activity is expressed as fold-repression values of normalized luciferase activity in test constructs compared to the CMV enhancer construct (ECMV). The full 1.2 kb sequence of the chicken 5’ HS4 insulator (cHS4) is used as positive control; a threshold line is indicated in red, corresponding to the enhancer-blocking activity of the cHS4 core sequence (II/III). As negative control a mutated version of II/III was used. Anova-Bonferroni multiple comparison test. Significant P<0.05. (**C**) Enhancer-blocking assay (EBA) of the *Tyr* 5’ element and its core sequence. (**D**) Enhancer-blocking assay in zebrafish embryos. Putative insulators are inserted at the *Asp*718 site franking a hindbrain enhancer and a somite promoter driving GFP. Insertion of the *Tyr* 3’ core element increases the fluorescence ratio between somites and hindbrain. Mutation of the CTCF binding site decreases the fluorescence ratio. Median test; n = 11-39. *Tyr* 3’ versus control, p<0.00726; *Tyr* 3’ mut versus control, p<0.135; *Tyr* 3’ versus *Tyr* 3’ mut, p<0.591. (**E**) Representative embryo injected with the empty pCAR48R vector. (**F**) Representative embryo injected with the *Tyr* 3’ core construct and (**G**) with the *Tyr* 3’ mutated core construct. The hindbrain location is indicated by a yellow box.

At this point, we reasoned that the sequence corresponding to strongest chromatin interaction emerged in our 3C assay – located 3’ upstream the *Tyr* gene – could also act as a chromatin boundary as well. We hypothesized that two boundary elements, at each end of the locus, would guarantee the strict cell-type specificity of *Tyr* expression and shield the locus from undesired interactions with neighbouring *Nox4* and *Grm5* genes (**Fig 3A**). Supporting our hypothesis, we observed that the *Tyr* 3’ region is occupied by CTCF, a protein that exert insulator activity, in several mouse tissues and cell lines (**Fig S3A**). Furthermore, a SINEB2 retrotransposon, a class of repeated mobile elements associated with insulator activity(36), was also found in this *Tyr* 3’ region (**Fig 3A**; **Fig S3B**), carrying a CTCF-binding motif (**Fig S3C**). We therefore interrogated the *Tyr* 3’ element for its enhancer-blocking activity using an enhancer-blocking assay (EBA). Indeed, the *Tyr* 3’ element functions as an insulator *in vitro*: the full-length, 2.5 kb *Tyr* 3’ element displayed the highest enhancer-blocking activity. Its core component, a 241 bp element containing the SINEB2 retrotransposon including a CTCF-binding site, retains more than 50% of the activity of the full-length counterpart in less than one tenth of the size. Upon mutating the CTCF binding site we could reveal the contribution of the SINEB2 element to the boundary activity, which was reduced, but not fully abolished (**Fig 3C**). This was expected, in the absence of the consensus CTCF binding motif(53).

In vivo, using an EBA in zebrafish embryos, where we tested the ability of a sequence to interrupt promoter-enhancer interactions, we confirmed the statistically significant insulator property of the *Tyr* 3’ core element (**Fig 3D**), that is capable of blocking enhancers *in vivo*. Likewise, upon introducing mutations at the CTCF site (**Fig S3D**), the resulting enhancer-blocking activity is reduced (**Figure 3D-F**) thereby proving that the boundary activity is, at least partially, CTCF-dependent. These results suggested that the *Tyr* expression domain is flanked by two elements capable of blocking enhancer activity *in vitro* and *in vivo*, by two chromosomal insulators

### The Tyr 3’ element is dispensable for Tyr expression

In order to functionally assess the role, in vivo, at the endogenous location, of the *Tyr* 3’ element, we used the CRISPR/Cas9 system to produce targeted chromosomal deletions involving this *Tyr* 3’ site. Guide RNAs were picked flanking a 2.8 Kb region encompassing the *Tyr* 3’ insulator, including the *Dpn*II fragment highlighted by the 3C assay and the sequences tested in the enhancer-blocking assays. We established two mouse lines carrying deletions of different sizes in homozygosis (TYRINS3#16 and TYRINS3#26; **Fig 4A-B**). Coat colour of TYRINS3 mice appeared indistinguishable from that of wild type pigmented mice (**Fig 4C**). The melanin contents from skin and eye extracts from both lines were comparable to those of wild-type animals (**Figure 4D-E**). Whole-mount retinae of *Tyr* 3’ animals were also indistinguishable from those of wild-type littermates (**Fig 4F-G**), and we did not observe delay in the appearance of eye pigmentation in developing embryos (**Fig S4A-B**) suggesting that perhaps the *Tyr* 3’ element might not be required for *Tyr* expression in melanocytes or RPE.

**Figure 4:**
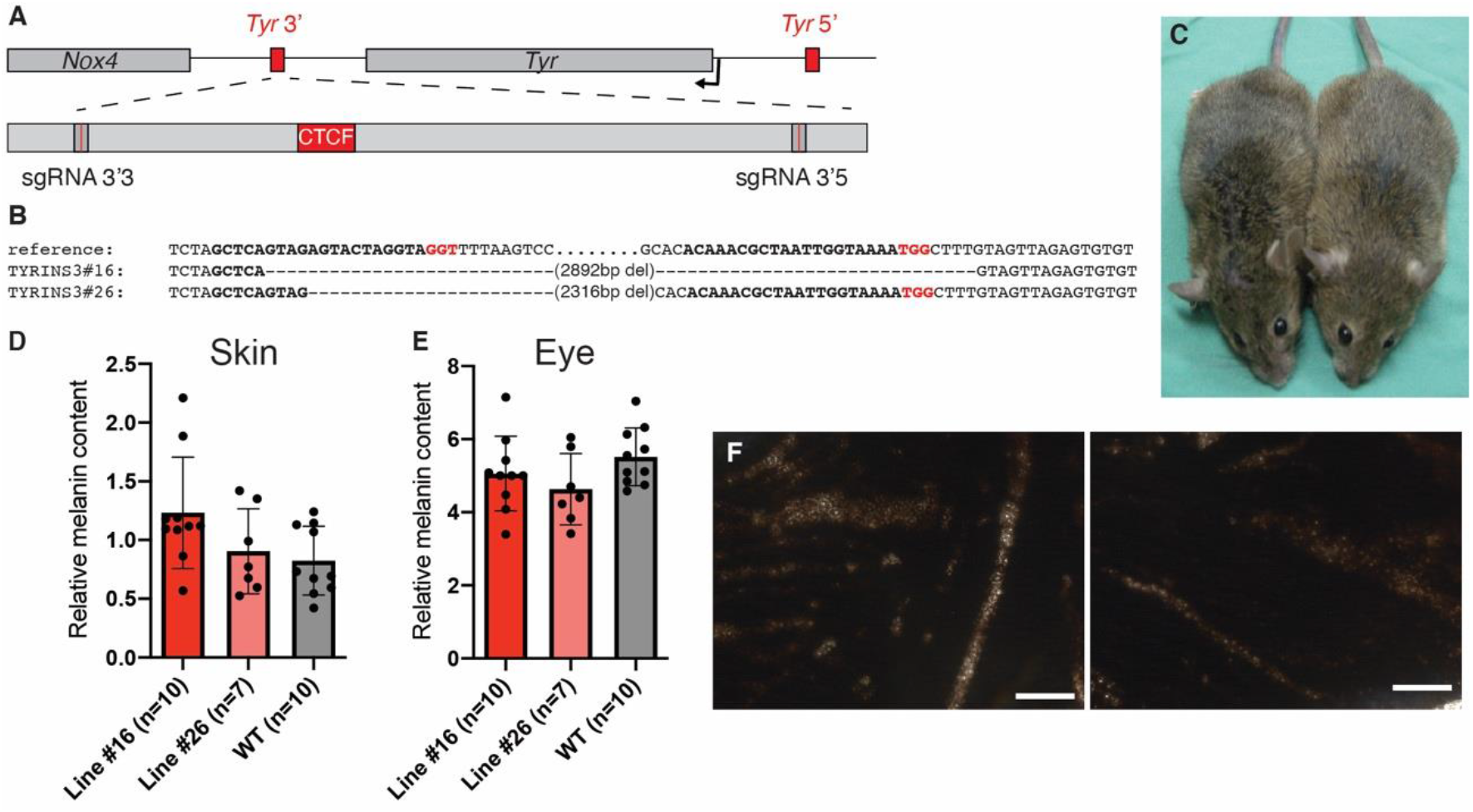
Deleting the *Tyr* 3’ element in mice does not alter the pattern *Tyr* gene expression. (**A**) Diagram of the *Tyr* locus illustrating the position of the sgRNAs used for deletion of the *Tyr* 3’ boundary. (**B**) Alignment of the deletion alleles in two mouse lines. Line TYRINS3#16 carries a 2892 bp deletion; line TYRINS3#26 carries a smaller deletion (2316 bp) that spares the sequence targeted by sgRNA 3’5. (**C**) The coat color of wild-type (left) and TYRINS3 homozygous (right) animals is indistinguishable. (**D**) Skin and (**E**) eye relative melanin content of TYRINS3#16, TYRINS3#26 homozygous and wild-type mice. (**F**) Whole mounts of retinae from wild-type (left) and TYRINS3 homozygous (right) animals.

### Inactivation of the Tyr boundaries perturbs the transcription of flanking genes in vivo

Insulators have been classically studied by means of reporter assays or by testing for copy-number dependent activity of integrated reporter genes(54). Such approaches limited our understanding of the actual impact of insulators on local gene expression and that of neighboring genes. Only recently, with the advent of genome engineering techniques(55, 56), the study of boundary elements at their endogenous context became accessible. This can be done by direct editing of CTCF-binding sites(57) or by introducing large rearrangements that disrupt topological chromatin domains(7). In this context, our mouse lines with genome-edited insulator sequences represent a unique opportunity to investigate the functional role of these two genomic insulators and their impact on the expression of *Tyr* and the adjacent *Nox4* and *Grm5* genes at their endogenous locus. For this purpose, we generated homozygous animal lacking the *Tyr* 3’ element (this work, TYRINS3 mice) or the *Tyr* 5’(34) (TYRINS5 mice), and quantified gene expression levels of *Tyr* and of the two flanking genes *Nox4* and *Grm5* in different tissues.

In the eye, loss of the *Tyr* 5’ element in TYRINS5(34) homozygous mice results in 50% reduction of *Tyr* expression. This finding is in line with previous histological analyses(34) and with its enhancer activity in melanocytes of the choroid (**Fig 2**). Deletion of the *Tyr* 3’ element in TYRINS3 homozygous mice does not affect *Tyr* expression in the eye. Once again, this finding is in agreement with eye melanin measurements (**Fig 4E**) and whole mount retina, that are indistinguishable from wild-type mice (**Fig 4F-G**). The expression of *Nox4* and *Grm5* remains unaltered in the eye of TYRINS3 and TYRINS5 homozygous mutant animals (**Fig 5A**), respectively.

**Figure 5:**
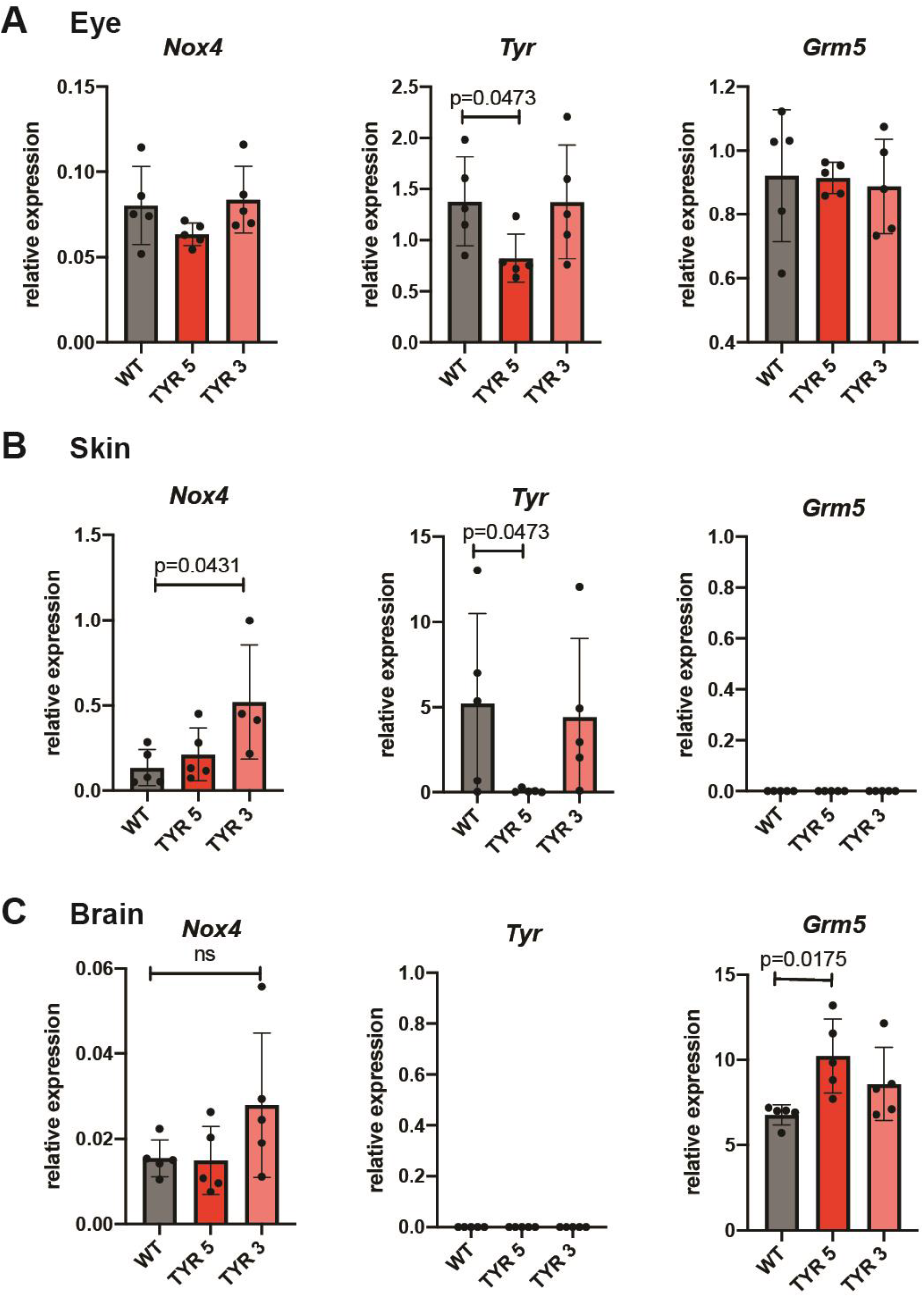
Inactivation of the *Tyr* boundaries perturbs the transcription of flanking genes in vivo. (**A**) Relative mRNA expression of *Tyr*, *Nox4* and *Grm5* in the eye of wild-type and homozygous TYRINS5 and TYRINS3 mice. (**B**) Relative mRNA expression in the skin and (**C**) brain of wild-type and homozygous TYRINS5 and TYRINS3 mice. Kruskal-Wallis test.

Loss of *Tyr* 5’ in TYRINS5 homozygous animals almost abolished *Tyr* expression in the skin, in agreement with their lighter coat colour and previous melanin measurements(34). Interestingly, deletion of *Tyr* 3’, the element which separates *Tyr* and *Nox4,* leads to perturbation in *Nox4* expression as evidenced by significant upregulation of *Nox4* transcripts in the skin of TYRINS3 homozygous mice (**Fig 5B**). This perturbation in the pattern and strength of *Nox4* expression in the absence of the *Tyr* 3’ element, further confirms its function as insulator (Fig 3B-D).

As previously reported(24), *Tyr* is not expressed in the the brain of wild-type animals and this pattern does not change upon deletion of the *Tyr* 3’ or *Tyr* 5’ elements. Conversely to *Tyr*, *Grm5* is a neuro-specific gene and is expressed in the brain of wild type mice and both *Tyr* 5’ and *Tyr* 3’ mutants. Once again, our findings clearly show that the loss of the boundary region can perturb expression of genes flanking the boundary. This is evidenced by the significant increase in the brain expression levels of *Grm5* upon deleting the *Tyr* 5’ element that separates the *Tyr* and *Grm5* loci, in TYRINS5 mice. Consistent with our model, deletion of the Tyr 3’ element separating *Tyr* and *Nox4* genes in TYRINS3 mice has no bearing on *Grm5* transcript levels (**Fig 5C**).

## DISCUSSION

In this work we describe the chromatin structure of the mouse *Tyr* locus by 3C in mouse melanoma (expressing *Tyr*) and fibroblast (non-expressing *Tyr*) cells. We have identified two chromatin loops: one involves the interaction between *Tyr* 5’ enhancer and the promoter, and correlates with *Tyr* expression; the other, detected in *Tyr* both type of cells, involves the structural protein CTCF, bound to a 3’ far upstream element.

The *Tyr* 5’ enhancer was initially identified in the classical “*chinchilla mottled*” *Tyr* mouse mutation (*Tyr*^*c-m*^)(28) and, thereafter, its pivotal role confirmed in mouse transgenesis experiments, through the rescue of pigmentation using both small(29) and large DNA constructs(32), or using reporter lines(50) in oocytes from albino mice. These experiments highlighted with high precision the location and the mechanism of the enhancer, that is transactivated by Sox10 and Mitf(50). USF1 transcription factor, usually found in chromatin insulators(52), was also reported to interact with this *Tyr* 5’ element(48), likely explaining the boundary activities described in mice(30, 34), flies(30) and zebrafish(39). Due to confounding factors, including overall transgene integrity, copy number and integration site, it remains challenging to assess up to what extent the enhancer is required to drive *Tyr* expression using classical transgenic mouse models, since even the smallest *Tyr* promoter tested in transgenic mice (270 base pairs in length) was sufficient to drive *Tyr* expression in both melanocytes and RPE cells(25).

More recently, thanks to advances in genome editing techniques(56, 58), we targeted the *Tyr* 5’ enhancer and boundary elements at the endogenous locus (TYRINS5 mice(34)). The phenotype of mice lacking the endogenous enhancer is less severe compared with what we had observed using engineered YACs(32, 33), where variegation was regularly reported, however highly reproducible across TYRINS5 mouse lines with similar deletions(34). Using the TYRINS5 mice, we determined that the *Tyr* 5’ element is absolutely required to drive *Tyr* expression in neural-crest derived melanocytes, where *Tyr* gene expression is almost undetectable upon deleting the *Tyr* 5’ element (**Fig 2B**, **5B**), but dispensable in the RPE (**Fig 2**). Mechanistically, this correlates well with differences in gene expression between melanocytes, that express Sox10, and cells of the RPE, that are Sox10 negative(50, 51). In fact, in melanocytes, Sox10 binding overlaps the core *Tyr* 5’ enhancer(46) (**Fig 2C**) that is enriched in Sox10 binding motifs (**Fig S2A**). A RPE-specific enhancer was identified using BAC transgenesis and transient LacZ reporter mouse lines(48, 49); however, the specificity and relevance of such distant enhancer (absent in some of the YAC *Tyr* transgenic mouse lines that were indistinguishable from wild-type pigmented mice, where DNA sequences upstream of the 5’ element had been deleted(32)) remain to be validated at the native chromatin context, and in relation with the *Tyr* promoter. These are ongoing experiments whose results will be discussed in a following study.

At the 3’ end of the locus, at the intergenic space between *Tyr* and *Nox4*, we identified a sequence that interacts strongly with the *Tyr* promoter in melanoma cells and fibroblasts (**Fig 1B**). This sequence contains a SINEB2 retrotransposon and it is bound by CTCF (**Fig S3**). These features are typically associated with insulators(36, 59), a class of regulatory elements able to block enhancer-promoter interactions and to partition chromatin in transcriptionally independent domains(60). Indeed, we could detect a strong insulator activity, in mammalian cells and in zebrafish embryos at this *Tyr* 3’ element comparable to that of the archetypic chicken cHS4 insulator, both CTCF- and SINEB2-dependant, (**Fig 3**). This further highlights the role of retrotransposons in the gene insulation(36, 37), in control of chromatin topology(61), and in genome evolution(62, 63). Deletion of this sequence in the mouse does not affect pigmentation or melanin levels in the skin or the eye of two independent mouse lines (**Fig 4**); *Tyr* mRNA level in the eye and skin does not depend of the *Tyr* 3’ insulator (**Fig 5**). However, when we measured the expression level of *Nox4*, located further upstream and beyond the *Tyr* 3’ boundary, we found that *Nox4* was overexpressed in the skin, suggesting that loss of the *Tyr* 3’ allowed *Tyr* regulatory elements to abnormally and ectopically transactivate *Nox4.* Similarly, we profiled gene expression in tissues of mice lacking the *Tyr* 5’ element. *Grm5*, the gene found 5’ upstream of *Tyr* and beyond the *Tyr* 5’, is overexpressed in the brain of animals lacking the *Tyr* 5’. We speculate that rearrangements in chromatin topology in the absence of the *Tyr* 5’ element expose the *Grm5* promoter to distal regulatory sequences. Several mouse models (**Fig 6A-B**) illustrated the relevance of regulatory regions at the mouse *Tyr* locus. Combining profiling of chromatin topology and functional study in the mouse, we devised a model for the locus: in melanocytes, the *Tyr* promoter engages two regulatory elements occupied by Sox10 and CTCF (**Fig 6C**). Disruption of the *Tyr* 3’ element, that normally interacts with the *Tyr* promoter and exert insulator activity, results in overexpression of the flanking gene *Nox4*, in the skin (**Fig 6D**). Perturbation of the *Tyr* 5’ element, that normally transactivate the *Tyr* promoter and functions as a boundary, results in loss of *Tyr* expression in gain in expression of the flanking *Grm5* (**Fig 6E**).

**Figure 6:**
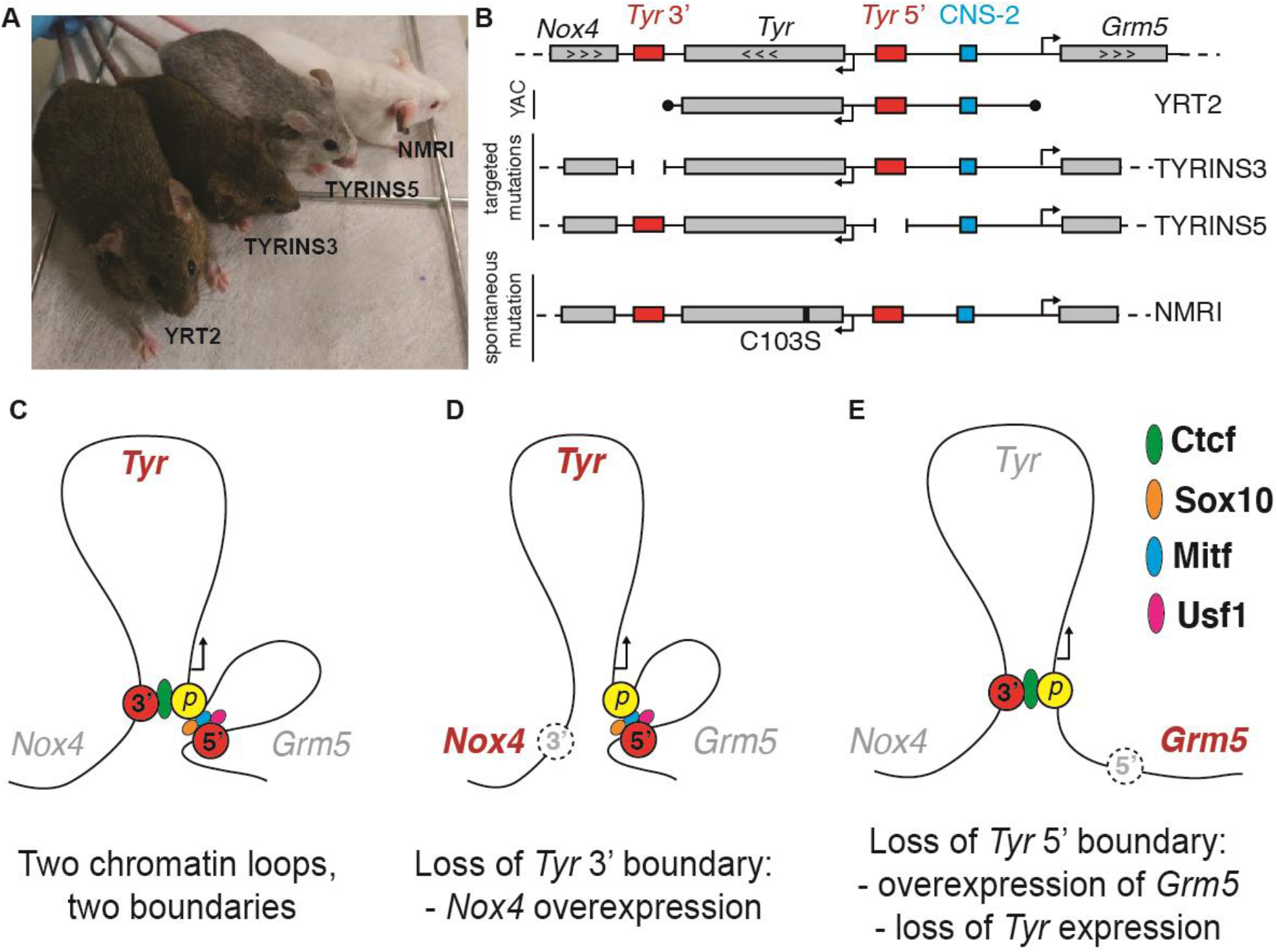
Model of the *Tyr* locus. (**A**) Representative individuals of mouse models for the *Tyr* locus: YRT2, TYRINS3, TYRINS5 and albino NMRI, along with (**B**) schematics of their genomic modifications. (**C**) When transcriptionally active, the *Tyr* locus is organized in two loops, mediated by CTCF at the 3’ boundary and likely by Sox10, Mitf and Usf1 binding at the 5’ boundary. (**D**) Inactivation of the *Tyr* 3’ boundary results in ectopic expression of *Nox4,* the gene located beyond the boundary. (**E**) Inactivation of the *Tyr* 5’ boundary causes loss of *Tyr* expression and overexpression of *Grm5,* past the boundary.

These data illustrate the role of enhancers and chromatin insulators in gene expression at the scale of a given locus and its adjacent genes. They also highlight how mutations at regulatory elements could alter the expression of genes in multiple tissues. For more than 20 years (1990(18)-2012) our ability to functionally and structurally interrogate the non-coding DNA sequences within the mouse *Tyr* locus was extremely limited. The extraordinarily abundance of DNA repetitive elements, including LINE1 retrotransposons in the vicinity of *Tyr*(30), rendered impossible any attempt to apply standard homologous recombination techniques in mouse embryonic stem (ES) cells. With the advent of the CRISPR/Cas9 techniques applied to mouse functional genomics(64), it has become possible dissecting the role of selected regulatory elements at their endogenous chromosomal location. This study highlights the relevance of investigating the non-coding genome in their native chromosomal context, and the advantage of using mouse models over cellular systems in the study of regulatory sequences.

## ACKNOWLEDGMENTS

The authors wish to thank the William J. Pavan and Stacie K. Loftus (NIH) for kindly providing melan-a ChIP-seq data. The authors wish to dedicate this work to the memory of Günther Schütz (1940-2020), who mentored L.M. between 1991 and 1995, and in whose laboratory mouse *Tyr* locus studies began.

