## supplementary material for "Boundary sequences flanking the mouse tyrosinase locus ensure faithful pattern of gene expression"

#### SUPPLEMENTARY TABLES AND FIGURES

**SUPPLEMENTARY TABLE**
**Table 1: Oligonucleotide sequences**

| | |
|---|---|
| **Primers used for 3C** | |
| **Anchor on *Tyr* promoter** | |
| AGGGGTTGCTGGAAAAGAAG | Tyr_promoter |
| TTTTCTTGACTTTGTGTCCCTATG | Tyr1 |
| TCTTAATGTTGGAAAGTGCAAAGT | Tyr2 |
| TCACAACACTGTCATACCATCTG | Tyr3 |
| TTGGACCTGTGCTGTGACTAA | Tyr5 |
| AAGGACGGAGTGGAGGTTG | Tyr6 |
| GGCAGGATGGGACTGAAGTA | Tyr11 |
| ATTTGTCTGGGGGCTCATAAC | Tyr12 |
| CACACAACTATCACTACTATCACCAC | Tyr13 |
| TGTTGAATCCCACCTTTACTCC | Tyr14 |
| CCATGCCCTGCTAAATGTGTA | Tyr15 |
| CCCAGACCCTTCCAAGTCAGTAT | Tyr16 |
| TGGAAAATGAGACACAACGAAG | Tyr17 |
| **Anchor on Tyr 5'** | |
| CACACAACTATCACTACTATCACCAC | Tyr_5' |
| TCACAACACTGTCATACCATCTG | Tyr3 |
| AAAGACACCATCCCTCCAAC | Tyr7 |
| TTCTCTTTTCTTTCGCACCA | Tyr8 |
| AGGGGTTGCTGGAAAAGAAG | Tyr9 |
| CCCAGACCCTTCCAAGTCAGTAT | Tyr16 |
| TACAGCAACACATTAGAACCAGA | Tyr17 |
| **Anchor on CNS-2** | |
| CCCAGACCCTTCCAAGTCAGTAT | Tyr_CNS2 |
| TCTCAAATCCCTCCTATCCAA | Tyr4 |
| TTGGACCTGTGCTGTGACTAA | Tyr5 |
| AAGGACGGAGTGGAGGTTG | Tyr6 |
| AGGCTGAGAGTATTTGATGTAAGAA | Tyr10 |
| CACACAACTATCACTACTATCACCAC | Tyr13 |
| TACAGCAACACATTAGAACCAGA | Tyr17 |
| ***Ercc3* locus** | |
| GTCTGTCTTTGTTGCTGAAGTATG | XBP1 |
| AAGTCCAGTGTGCTGAGGTATT | XBP2 |

| Primers used for cloning EBA vectors | |
|---|---|
| CAGCTCGAGACAGAAATGGCCCCACCTAT | XhoI_tyr3'F (2.5kb) |
| CAGCTCGAGTGCATTTGAACTTGACCTACTGA | XhoI_tyr3'R (2.5kb) |
| CAGCTGCAGACAGAAATGGCCCCACCTAT | PstI_tyr3'F (2.5kb) |
| CAGCTGCAGTGCATTTGAACTTGACCTACTGA | PstI_tyr3'R (2.5kb) |
| CAGCTGCAGCCAGGTGAGGGGTGTGTTTA | XhoI_tyr3'F (241bp) |
| CAGCTGCAGGAAGTGTTTATTGACAATGTG | XhoI_tyr3'R (241bp) |
| CAGCTCGAGCCAGGTGAGGGGTGTGTTTA | PstI_tyr3'F (241bp) |
| CAGCTCGAGGAAGTGTTTATTGACAATGTG | PstI_tyr3'R (241bp) |
| TGTCTTCAGACACTCGAGAATAGAGCGCCAGATCTTGTTA | 3'CTCFmutF |
| TAACAAGATCTGGCGCTCTATTCTCGAGTGTCTGAAGACA | 3'CTCFmutR |
| **primers for cloning of sgRNAs** | |
| ACACCAGCTCAGTAGAGTACTAGGTG | Tyr3'3gRNAFw |
| AAAACACCTAGTACTCTACTGAGCTG | Tyr3'3gRNARv |
| ACACCACAAACGCTAATTGGTAAAAG | Tyr3'5gRNAFw |
| AAAACTTTTACCAATTAGCGTTTGTG | Tyr3'5gRNARv |
| **Primers for mouse genotyping** | |
| CAACCAGGCTTTCATCAGAAT | Tyr5_delF |
| TTTTCTCTGTATCATGATTGCCTA | Tyr5_delR |
| TCTGTGCATGGTATACAACAGGG | Tyr3_delF |
| GTGCATTAAAGGAAGCCCAATGA | Tyr3_delR |

#### Supplementary Figure 1

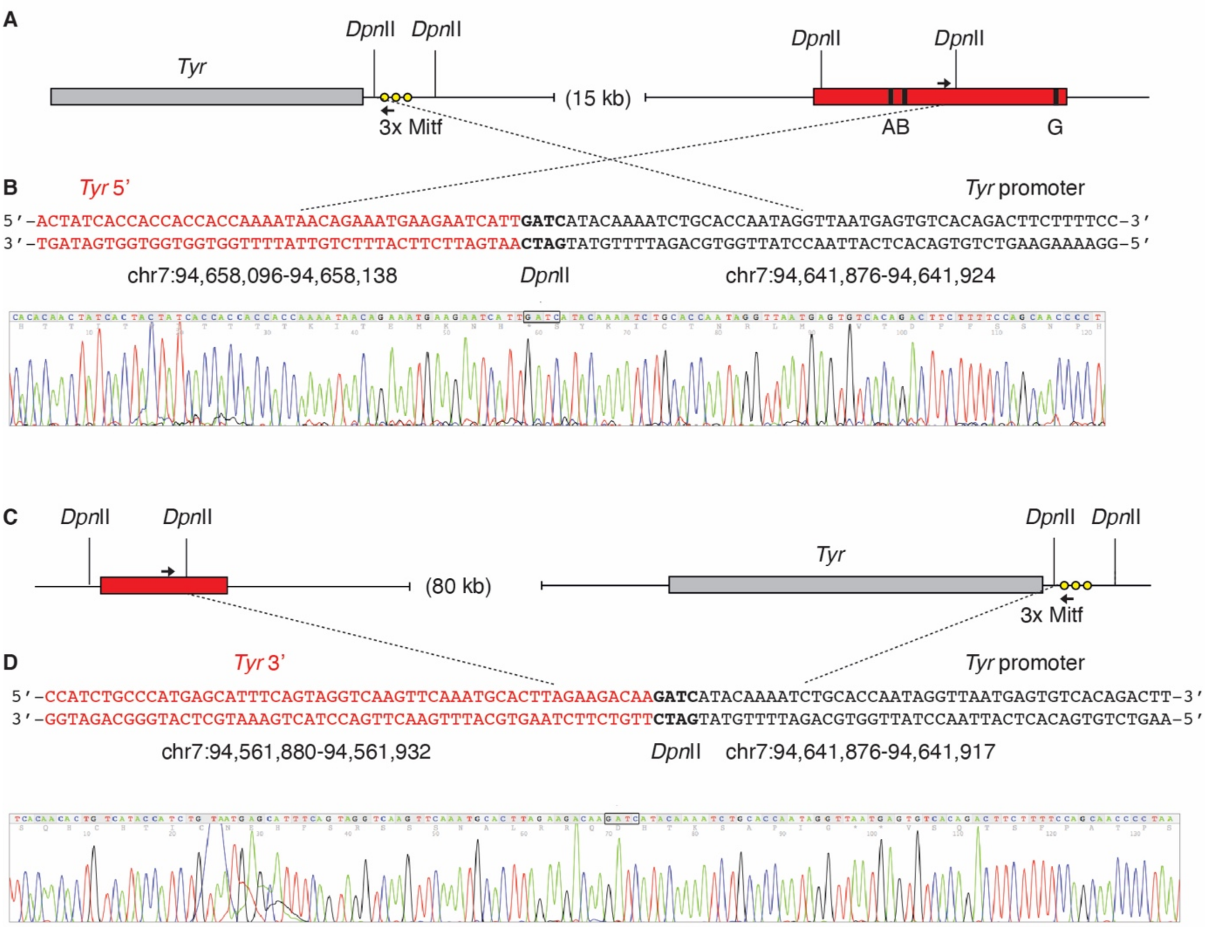

### Supplementary Figure 2

**A**

chr7-94,656,561-94,656,970 (mm9); **Sox10**, **Usf1**, **(Mitf)**

```
ATAACAGGAAAATAAACAAGCCACATTCCTGCTTTGTAGAAGTTCACAATCTGAATAGGAATATAAAAAGGTGATGGTGTGGCAATTTCAGGACTCTTTATTGGGTACA
TATTGTTAATAATATTGTGGTTTGCCAGGACCCCAGCAGGAAGCAGCTGACACACTCAAGCTGGTTCTTTGAACAGAGTTTAAGAAAGAAGCTATTTACAATGTTGGGG
                                                                          B box                     A box
GTGGAATTTAAGAAGAAAGGACAGTGGATAATGAAAAGGATACTCAGAACAACTGCTGGAACACAGACATCACTGTCTTGACAA(AGTCAGCTGGC)AGGAACTATGACCT
TGGAGAAGGATATAAGCAACTTTCTTGTTATGTGGCAAAGAGGAAGTTGCAGACCTTTGACTTCACTCTCCTTCTGCCTCTAA
```

**B**

**TRANSFAC**

| TF | KO mouse | pigmentation |
|---|---|---|
| Ets | yes | normal |
| Cdx2 | lethal | n.a. |
| Runx2 | yes | normal |
| Sry | yes | normal |
| Nfatc2 | yes | normal |
| Nr5a2 | lethal | n.a |
| Myb | yes | normal |
| Ttf1 | n.a. | n.a |
| Bptf | lethal | n.a. |
| **Lef1** | **yes** | **aberrant melanocytes with no melanin** |

**UNIPROBE**

| TF | KO mouse | pigmentation |
|---|---|---|
| Lmx1b | yes | normal |
| Sry | yes | normal |
| Pou4f3 | yes | normal |
| **Lmx1a** | **yes** | **white spotted** |
| Arid1a | yes | normal |
| Sox7 | lethal | n.a. |
| Sox12 | yes | normal |

#### Supplementary Figure 3

**A**

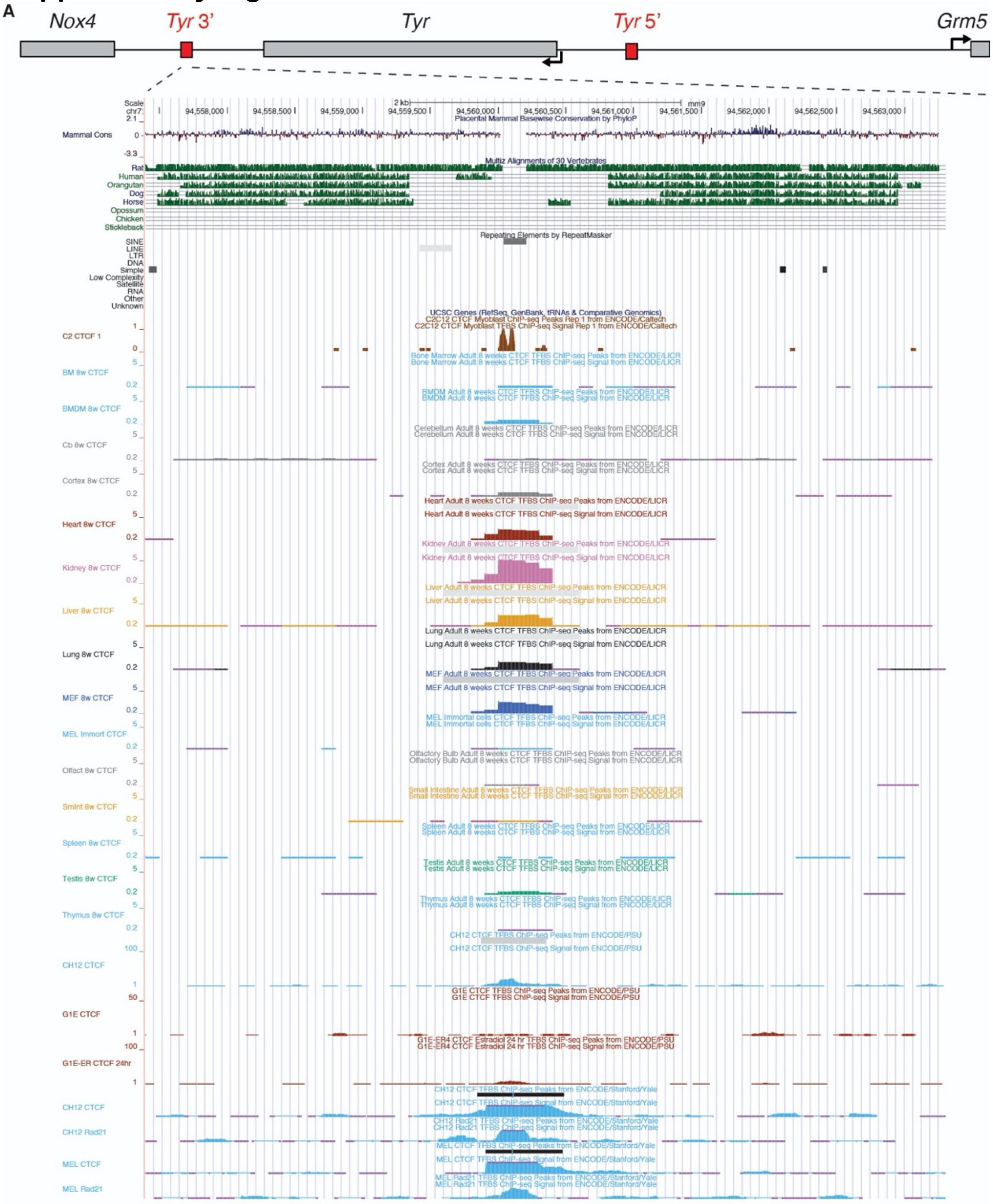

**B** GAAGTGTTTATTGACAATGTGACATTTTGTTTTG**TTTTGTTTTTTAAAGATTTATTTATTTATTATATATGTAAGTTC**
**ACTGTAGCTGTCTTCAGACACTCCAGAAGAGGGCGCCAGATCTTGTTACGGATGGTTGTGAGCCCCCATGTGGTTG**
**ACCTTCGGAAGAGCAGTCAGTGCTCTTAGCTGCTGAGCCATCTCTCCAGCCC**TGACATTGTGATATTTTTAAACAC
ACCCCTCACCTGG

chr7:94560004-94560244 (minus strand)

**C** GAAGTGTTTATTGACAATGTGACATTTTGTTTTGTTTTGTTTTTTAAAGATTTATTTATTTATTATATATGTAAGTTC
ACTGTAGCTGTCTTCAGAC**ACTCCAGAAGAGGGCGCCAG**ATCTTGTTACGGATGGTTGTGAGCCCCCATGTGGTTG
ACCTTCGGAAGAGCAGTCAGTGCTCTTAGCTGCTGAGCCATCTCTCCAGCCCTGACATTGTGATATTTTTAAACAC
ACCCCTCACCTGG

**D** 5′-ACTCCAGAAGAGGGCGCCAG-3′ wt

5′-ACTC**G**AGAA**T**AG**A**GCGCCAG-3′ mutated

**Supplementary Figure 4**

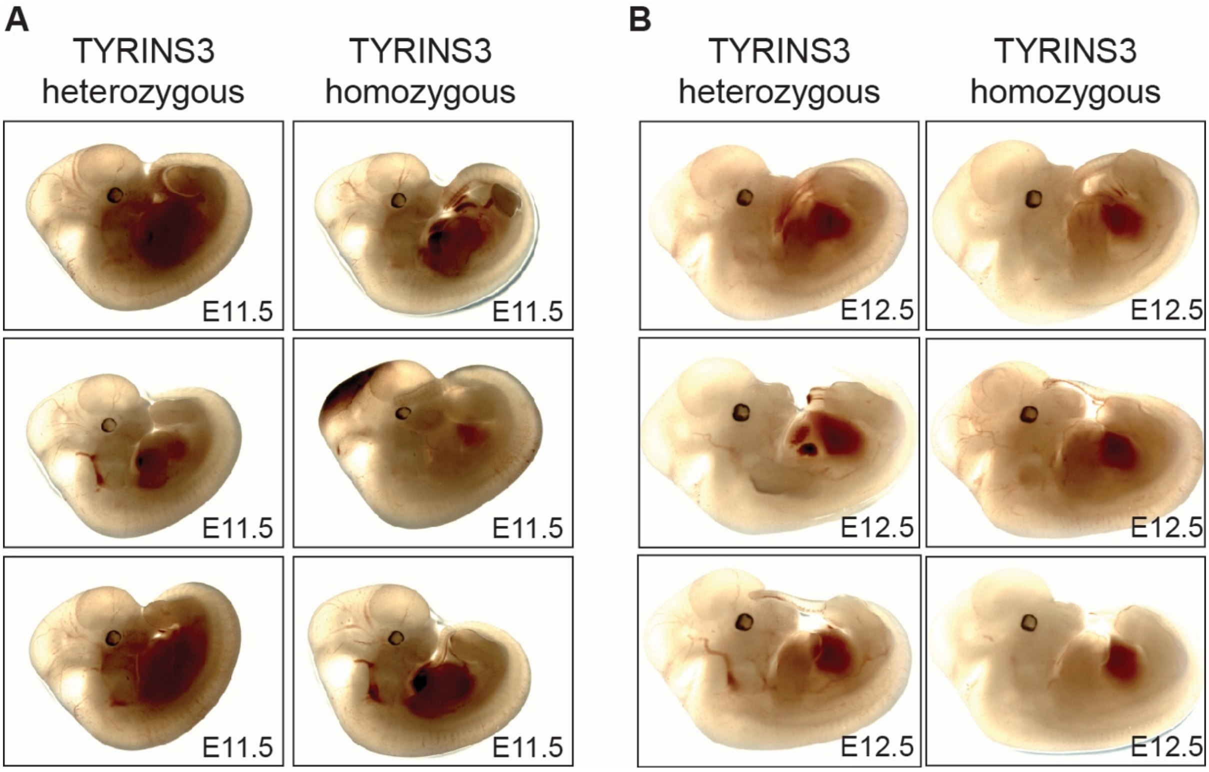

#### SUPPLEMENTARY FIGURE LEGENDS

**Figure S1: Chromosome conformation of the mouse *Tyr* locus.**
(**A**) Diagram of the DpnII restriction sites at the *Tyr* promoter and *Tyr* 5' element. MITF binding sites at the *Tyr* promoter are indicated as yellow circles. A, B and G boxes are highlighted in black. 3C primers are depicted as black arrows (**B**) Sanger sequencing of chimeric ligation products resulting by proximity ligation of DpnII fragments at the *Tyr* promoter and *Tyr* 5' element. (**C**) Diagram of the DpnII restriction sites at the *Tyr* promoter and *Tyr* 3' element. (**D**) Sanger sequencing of chimeric ligation products resulting by proximity ligation of DpnII fragments at the *Tyr* 3' element and *Tyr* promoter. Genomic coordinates are indicated (mm9)

**Figure S2: The *Tyr* 5' element contains a melanocyte-specific *Tyr* enhancer**
(**A**) Sequence of the *Tyr* 5' core enhancer. Sox10 binding motifs are highlighted in red; Usf1 motif is highlighted in green, with overlapping Mitf motif between brackets. A and B box sequences are boxed (**B**) Additional transcription factor binding motif predictions using TRANSFAC and UNIPROBE.

**Figure S3: The *Tyr* locus is flanked by chromatin boundaries**
(**A**) Genomic view of the *Tyr* 3' element. Mammalian sequence conservation and repeat DNA tracks from USCS Genome Browser. CTCF occupancy in mouse cell lines and tissues from ENCODE (**B**) Sequence of the *Tyr* 3' core element; the SINEB2 sequence is highlighted in red. (**C**) The CTCF binding motif is highlighted in red. (**D**) CTCF binding motif compared with its mutagenized version used in zebrafish.

**Figure S4: Deleting the *Tyr* 3' element in mice does not alter the pattern *Tyr* gene expression.** (**A**) Heterozygous and homozygous TYRINS3 E11.5 embryos (**B**) Heterozygous and homozygous TYRINS3 E12.5 embryos.
